# Binding of Small Molecule Inhibitors to RNA Polymerase-Spt5 Complex Impacts RNA and DNA Stability

**DOI:** 10.1101/2023.09.12.557491

**Authors:** Adan Gallardo, Bercem Dutagaci

## Abstract

Spt5 is an elongation factor that associates with RNA polymerase II (Pol II) during transcription and has important functions in promoter-proximal pausing and elongation processivity. Spt5 was also recognized for its roles in the transcription of expanded-repeat genes that are related to neurodegenerative diseases. Recently, a set of Spt5-Pol II small molecule inhibitors (SPIs) were reported, which selectively inhibit mutant huntingtin gene transcription. Inhibition mechanisms as well as interaction sites of these SPIs with Pol II and Spt5 are not entirely known. In this study, we predicted the binding sites of three selected SPIs at the Pol II-Spt5 interface by docking and molecular dynamics simulations. Two molecules out of three demonstrated strong binding with Spt5 and Pol II, while the other molecule was more loosely bound and sampled multiple binding sites. Strongly bound SPIs indirectly affected RNA and DNA dynamics at the exit site as DNA became more flexible while RNA was stabilized by increased interactions with Spt5. Our results suggest that the transcription inhibition mechanism induced by SPIs can be related to Spt5-nucleic acid interactions, which were altered to some extent with strong binding of SPIs.

## INTRODUCTION

RNA polymerase II (Pol II) is a vital protein known for its fundamental role in RNA synthesis, which takes place in three main stages that are initiation, elongation, and termination [1,2]. The transcription elongation stage starts with promoter-proximal pausing that causes Pol II to escape from the promoter site to maintain RNA elongation. During the elongation stage, several transcription elongation factors associate with Pol II and collectively regulate the transcription process [3]. Spt5 is one of the elongation factors that associates with Spt4 to form 5,6-dichloro-1-β-D-ribofuranosylbenzimidazole (DRB) sensitivity-inducing factor (DSIF) [4]. DSIF is known to have many roles in transcription including in promoter-pausing [5,6], processivity of transcription [7,8], nucleosomal access [9-11], and is recently recognized for its role in selective regulation of transcription of repetitive genes [12-15]. Although several studies in structural biology [6,7,10,16-22], biochemistry, and genetics [9,23-30] provided substantial data and valuable insights into DSIF function, understanding the mechanisms, in which DSIF plays a role in these transcription events is still limited. Structural studies showed that Spt5 interacts with Pol II upstream and stabilizes DNA and RNA at the exit tunnels [7,16]. Our previous study also shows that KOW1 and KOW4 domains stayed strongly in contact with the upstream DNA and nascent RNA, respectively, during the MD simulations [31]. These studies suggest strong interactions of DSIF with DNA and RNA, and these interactions could be directly related to the functional roles of DSIF in transcription elongation.

The role of DSIF in the transcription of genes with multiple repeats was associated with several neurodegenerative diseases [12-15]. Studies showed that reducing the expression of DSIF orthologs selectively inhibits the transcription of repetitive genes [12-15], which suggests a crucial role of DSIF in the transcription of such multi-repeat genes. Therefore, interference with DSIF function may have therapeutic consequences by decreasing pathological outcomes of neurodegenerative diseases such as Huntington’s disease [32], spinocerebellar ataxia type 36 [33], amyotrophic lateral sclerosis (ALS) and frontotemporal dementia [34,35], which are all known to be related to expanded gene mutations. Following these studies on the effects of DSIF in multi-repeat gene transcription, a recent study suggested a novel therapeutic approach for combating Huntington’s disease by introducing Spt5-Pol II small molecule inhibitors (SPIs), which selectively inhibit transcription of the mutant huntingtin (Htt) gene, which has extended CAG repeats, by reproducing the DSIF knockdown effects [36]. They observed inhibitory effects of SPIs on the transcription of the mutant Htt gene as well as inflammatory genes, while the mechanisms of how these molecules interfere with DSIF function and how they interact with the Pol II complex are open questions. Fluorescence intensity measurements on selected two SPIs suggest different binding patterns as one of the SPIs interacts with mostly Pol II Rpb1 subunit, while the other one seems to have interactions with both Rpb1 and Spt5. Although the earlier study provides some insight into their binding, the structural details of the binding sites are not fully resolved, yet crucial to understanding the molecular mechanisms of actions for these SPIs in transcription inhibition.

In this study, we predicted the binding sites of SPIs at the Pol II-DSIF complex and further elucidated interactions between the SPIs and the residues of Pol II and Spt5. We selected three SPIs that are known to selectively inhibit the mutant Htt gene transcription and performed docking calculations to determine their initial binding poses. Then, we performed molecular dynamics (MD) simulations to obtain binding strengths, interacting residues, and the impact of these interactions on the dynamics of the Pol II-DSIF complex. We observed that two SPIs are strongly bound to the initial binding sites predicted by docking at the Pol II-Spt5 interface, whereas one SPI demonstrated multiple binding sites. Notably, we observed that the strong binding of the formers altered Spt5-nucleic acid interactions that further impacted the stability of DNA and RNA at the exit site while weakly binding SPI had less impact on the interactions and dynamics of the nucleic acids.

## METHODS

### Initial structures of SPIs

We selected three molecules that inhibit the transcription of the mutant Htt gene from the previously published list of SPIs.[36] The SPIs I, II, and III correspond to 18, 21, and 86 in the previous paper [36] and the structures were shown in Fig. 1. The initial coordinates were generated using the ChemDraw program online version (19.0.0-CDJS-19.0.x.9+da9bec968) and the Avogadro program version 1.2.0. We first generated 2D structures using ChemDraw and then converted 2D structures to 3D coordinates using the Avogadro program. Finally, the conformations were optimized using Gaussian09 [37] with MP2 geometry optimization method and 6-31G(d) basis. In addition, we used the B3LYP algorithm to generate optimized conformations and compared the results with MP2 models. Fig. S1 shows that both algorithms provided very similar conformations. We note that SPI-I has a chiral center and we performed simulations of both S- and R-enantiomers (S-I and R-I). We presented the results of R-I in the main text as this enantiomer has stronger binding to the complex and more impact on the dynamics of nucleic acids while S-I manifested more loosely binding and has a smaller impact on the DNA dynamics (Fig. S2).

**Fig. 1.**
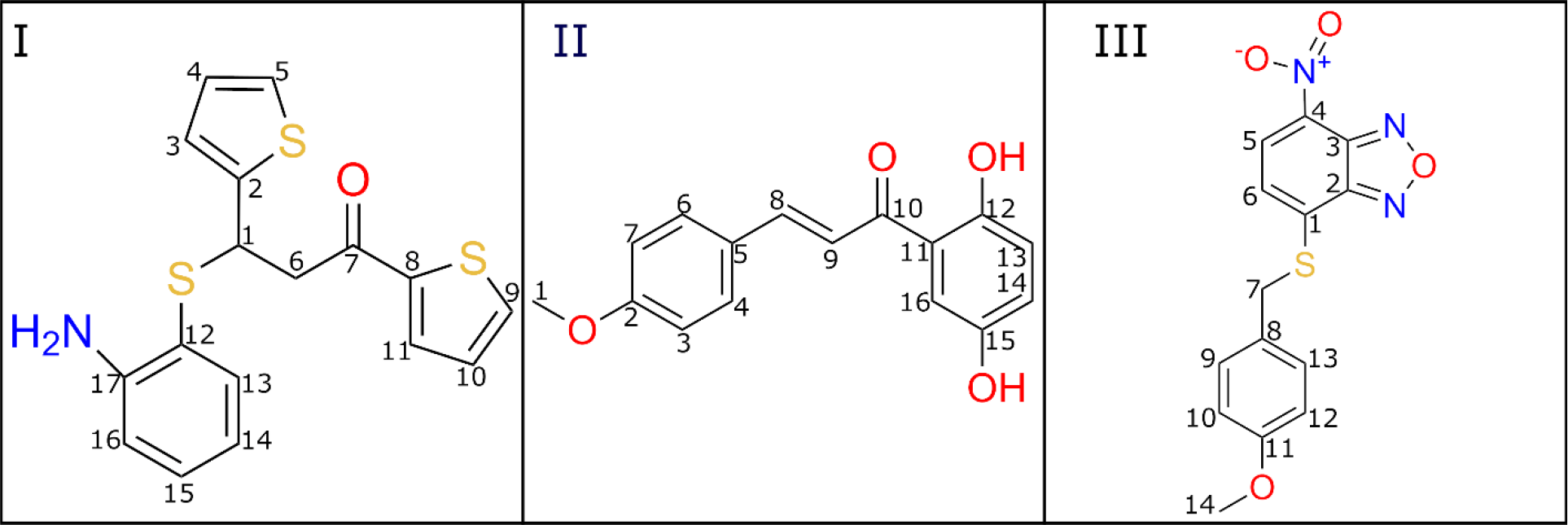
The chemical structures of SPIs used in this study.

### Docking of SPIs on the Pol II-DSIF complex

We used a recently published cryogenic electron microscopy (cryo-EM) structure of the Pol II elongation complex with the PDB ID of 5OIK [16]. We performed docking using the AutoDock-GPU [38], a graphic processing unit (GPU) accelerated version of the AutoDock program version 4.2.6 [39], and prepared input files for coordinates and grid maps using AutoDockTools [39]. Kollman [40] and Gasteiger [41] charges were used for protein and ligand partial charges, respectively. Only polar hydrogens were added to the protein. For docking, a grid with a size of 60 Å at each direction with a grid spacing of 0.375 Å was generated to cover the interface between the Pol II Rpb1 subunit and the Spt5 factor as this region was suggested as the binding site by fluorescence intensity measurements [36]. Docking was performed using the default parameters of AutoDock-GPU except for the number of genetic algorithm runs, which was set to 1000. Briefly, we used the Lamarckian genetic algorithm with the ADADELTA gradient-based local search method with a 100 % of local search rate, the rates of gene mutation and crossover were 2 % and 80 %, respectively, and the maximum number of local search iterations was 300. We performed all the docking runs on the OpenCL platform using NVIDIA Quadro RTX 6000 GPUs. The best binding conformation with a minimum total binding energy among all the docking conformations was used for each ligand for MD simulations.

### MD simulations

We performed MD simulations of protein-ligand complexes for three SPIs as well as for the apoprotein (apo). All the systems were prepared using the MMTSB toolset [42] and CHARMM software version 45b2 [43]. The protein complexes were solvated in a cubic box with a 10 Å cutoff from each side of the box. We added Na^+^ ions to neutralize the systems. The systems have 177 Å box sixes that consist of around 560,000 atoms. We used the CHARMM c36 and c36m force fields [44-46] for nucleic acids and proteins, respectively, and CGenFF [47] for SPIs by generating topology and parameter files using the CHARMM-GUI server [48,49]. The CHARMM-modified TIP3P water model was used for water molecules [50]. The masses in the force fields were modified to repartition the masses of the heavy atoms that are attached to H atoms to increase the mass of H to 3 a.m.u. as suggested earlier [51]. This modification increases the sampling of the simulations by allowing the use of a 4 fs time step without disturbing the system [51]. The energy of each system was minimized using 5000 steps with an energy tolerance of 100 kJ/mol. Systems were equilibrated for around 1.6 ns while the temperature was increased from 100 to 300 K and the time step was increased from 1 to 4 fs. During the equilibration, backbone and side chain heavy atoms of proteins were restrained using harmonic force constants of 400 and 40 kJ/mol/nm^2^, respectively. Ligands were not restrained during the equilibration steps. The analyses of root mean square deviation (RMSD) and densities show that the ligand conformations were slightly changed and their positions along the complex were unaltered during the equilibration (Fig. S3). After the equilibration, we performed MD simulations for 200 ns. Equilibration and MD simulations were repeated for three replicates for each system. For the long-range interactions, periodic boundary conditions were used with the particle mesh Ewald algorithm. Lennard-Jones interactions were switched between 1.0 to 1.2 nm. Langevin thermostat was used at 303.15 K and with 1 ps^-1^ friction coefficient. All the production runs were performed using 4 fs time step and trajectories were saved at every 40 ps. Simulations were performed using the OpenMM program [52] on GPU machines.

### Analysis of the simulations

Simulations were analyzed for RMSD, root mean square fluctuation (RMSF), distance, and contact maps, the relative center of mass of ligands, and ligand densities using the MDAnalysis package [53] and MMTSB toolset [42]. RMSD and RMSF of the proteins and nucleic acids were calculated for C_α_ and P atoms, respectively, after aligning C_α_ and P atoms of the frames onto a reference structure. For the RMSD calculations, the reference structure was the initial structure, while for the RMSF calculations, frames were aligned to the average structure over the trajectories. RMSD values of ligands were calculated for heavy atoms after superimposing the frames to the initial structure. Distance maps were calculated as the average minimum distance for each residue pair along the simulations. A principal component analysis (PCA) was applied to the distances between KOW4 and RNA for the apo-complex and the complex with SPI-I. For this analysis, we only included the pairs that have average distances smaller than 10 Å along the simulations of the complex with SPI-I. Contact maps were calculated for the minimum distance of ligand to each residue of Spt5 and Rpb1, and 5 Å used as the cutoff for the contact. The conformations of SPIs obtained by MD simulations were clustered based on RMSD values with respect to the initial structures using Kmeans clustering algorithm implemented in Scikit-learn Python module [54]. The conformations that are closest to the cluster centers were taken as the central structures for the corresponding clusters.

To evaluate the location of the ligands along the trajectories we calculated the relative center of mass of the ligands and ligand densities. The relative center of mass was calculated as the center of mass of the ligand with respect to the center of mass of the protein complex throughout the trajectory. Then, a PCA was applied to the relative center of mass of the ligand to obtain the corresponding positions of the ligand in a two-dimensional plot. After that, we performed the weighted histogram analysis method (WHAM) [55] to obtain the most probable positions of ligands along the protein complex. Ligand densities were calculated by applying a three-dimensional grid with 1 Å resolution on the trajectories that are aligned to the initial structures.

Binding energies from MD simulations were calculated by the molecular-mechanics and generalized Born with surface area (MM/GBSA) method using the CHARMM software and MMTSB toolset. To calculate binding energies, we first calculated the energies of protein and ligand alone as well as the energies of the protein-ligand complex. Then, binding energies were calculated by subtracting the sum of the protein and ligand energies from the energies of protein-ligand complex. We calculated binding energies of the SPIs to Pol II-DSIF complex and the contributions from the electrostatic and, van der Waals (vdW) interactions, as well as the electrostatic and hydrophobic solvation free energies, which were calculated by generalized Born (GB) [56] and atomic solvation parameter (ASP) energy terms [57], respectively, implemented in the Generalized Born Molecular Volume (GBMV) module in CHARMM using the analytical method [58]. ASP energy consists of the cavity term only which is calculated from the solvent accessible surface area (SASA) and surface tension parameter that is set to 0.015 kcal/mol/Å^2^. GB term used the generalized Born equation with dielectric constant of 80 for the solvent and 1 for the solute. For MM/GBSA calculations, we extracted frames every ns up to a total of 600 frames over three replicate trajectories for each system and perform a short minimization for each frame (50 steps of steepest descent and 50 steps of adopted basis Newton-Raphson) before the energy calculations.

## RESULTS

SPIs were reported to interfere with the DSIF function in the mutant Htt gene transcription. The previous study [36] predicted that SPI I interacts with the coiled-coil (CC) domain of the Rpb1 subunit and Spt5 elongation factor, while II showed interactions only with Rpb1. However, their exact binding sites were not predicted in the earlier study, therefore, in this work, we studied the binding of the three SPIs (Fig. 1) to the Pol II-DSIF complex. Below, we presented binding sites obtained by docking and MD simulations, and the impacts of the binding to the protein complex and nucleic acids.

### Docking predicted binding of the SPIs to the Pol II-Spt5 interface

We applied docking of the three SPIs to the Pol II-DSIF complex using a grid that covers interface of Spt5 and Rpb1 domain of Pol II in agreement with the previous experimental study [36]. The most favorable docking poses with the minimum energies for all the molecules were similar (Fig. S4). The docking positions are close to the CC domain of the Rpb1 subunit of Pol II for all the SPIs. Fig. 2 shows docking positions for the SPIs with highlighted interaction sites within 5 Å distance. SPI-I showed interactions with polar and charged residues (R291, R292, N287, N288) as well as with the nonpolar (I88) and aromatic (Y43) residues of the Rpb1 subunit of Pol II. The only nearby residue from Spt5 was E384, which may form H-bonds with the amine group of the SPI-I. II has interaction sites similar to I, but it is located closer to K42 and Q295, and lacks the interaction site with Y43 compared to I. III is supported by hydrophobic residues of Y43 and I88, and polar residue R292, while it is located relatively far from E384 of Spt5 compared to I and II. In addition, docking energies are provided in Table 1. II and III have lower energies compared to I, suggesting that they are more strongly bound to the docking site. However, our MD simulations suggest that II is more loosely bound compared to I and III, which we discussed below in more detail.

**Table 1.** Binding energies from docking and MM/GBSA calculations.

| SPI | Docking | MM/GBSA |  |  |  |  |
| --- | --- | --- | --- | --- | --- | --- |
|  | Energy | Total | Electrostatic | vdW | GB | ASP |
| I | -5.84 | -26.41±0.21 | -19.39±0.98 | -32.41±0.25 | 37.26±1.01 | -11.88±0.06 |
| II | -6.95 | -16.93±0.54 | -17.78±1.17 | -19.08±0.54 | 28.47±1.23 | -8.55±0.17 |
| III | -7.13 | -30.53±0.27 | -23.16±1.05 | -34.63±0.18 | 39.17±1.04 | -11.90±0.04 |

**Fig. 2.**
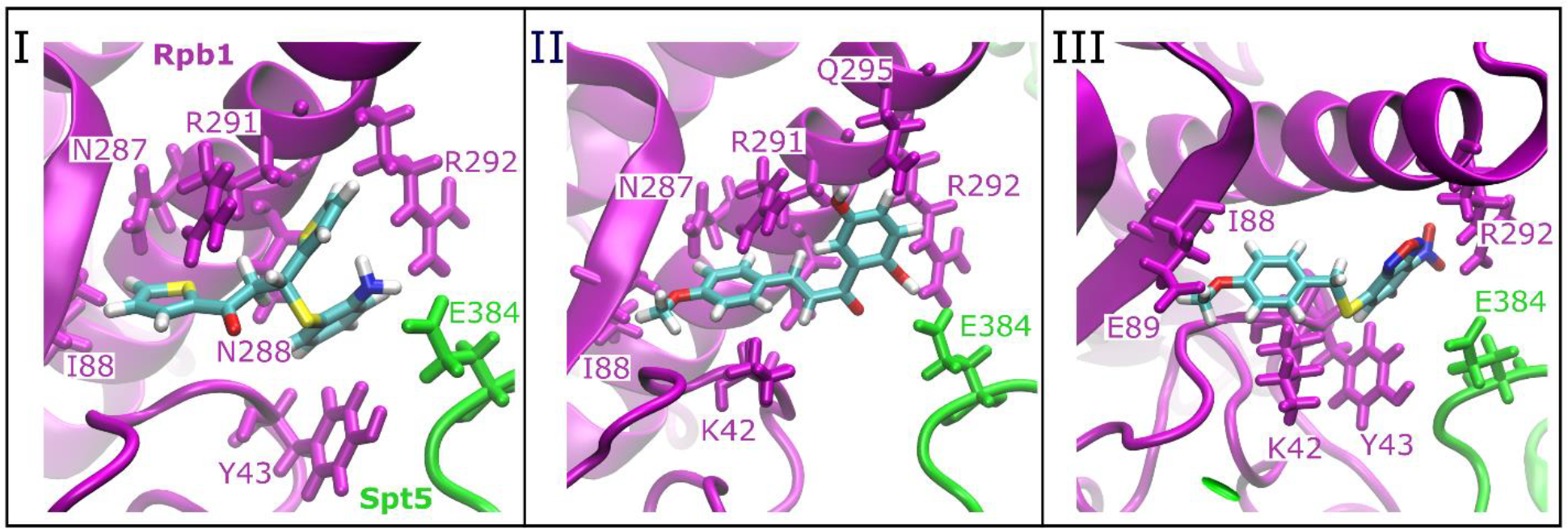
Docking positions for SPIs I, II and III at the Pol II-DSIF complex and the nearby residues of Rpb1 and Spt5 that are within 5 Å of the SPIs. The color code for the Pol II-DSIF complex is as follows: Rpb1 is in magenta, and Spt5 is in green; SPIs are shown in colors coded by atom name: C is cyan, H is white, O is red, N is blue, and S is yellow.

Docking energies of the minimum energy poses are shown in kcal/mol. MM/GBSA energies were calculated from the MD simulations trajectories and reported in kcal/mol. vdW, GB and ASP stand for the van der Waals, generalized Born and atomic solvation parameter energy terms, respectively.

### Two SPIs manifested strong binding to the Pol II-DSIF complex

We performed MD simulations of Pol II-DSIF complexes with SPIs at the minimum energy docking poses as the initial structures. Fig. S5 shows the RMSD plots of the SPIs during the three replicates of the simulations. SPI-I shows higher flexibility due to the rotation of the aromatic rings around the single bonds. SPI-II and III have fewer possibilities of rotation, as SPI-I has seven rotatable bonds while II and III have six and five rotatable bonds, respectively, which make them less flexible. In addition, SPI-II has an extended resonance that includes the double bonds of the C2-C7 aromatic chain and C8-C9 double bond (see Fig. 1). This may further decrease the flexibility of SPI-II. To classify the conformations from the MD simulations, we clustered them based on the RMSD values and showed the conformations at the cluster centers for each SPI (Fig. S6). We observed that the initial conformations were preserved to some extent while rotations of aromatic rings, and methyl and nitrogen dioxide groups are observed. In addition, SPI-I and SPI-III formed more bending structures compared to the initial optimized structures (Fig. S1, S6). The docking poses for the three SPIs were at the interface between the CC domain of Rpb1 and Spt5 (Fig. S4). Fig. 3a shows the densities of SPIs along the MD simulation trajectories, which show that for all three SPIs the initial docking site is one of the major binding sites. I and III strongly bound to the CC-Spt5 interface, while II sampled multiple binding sites. Fig. 3b-c shows that the major binding sites of SPI I and III are very similar to each other. These binding sites consist of the CC domain of Rpb1 and spatially nearby residues, and mostly NGN, KOW1 and KOW2-3 domains of Spt5. For II, the contacts are more scattered, nevertheless around similar regions, which are the CC domain of Rpb1, and NGN, KOW1, and KOW2-3 domains of Spt5. In addition, binding energies were calculated by the MM/GBSA method (Table 1). SPIs I and III bind more favorably compared to II consistent with the strong binding of I and III observed in Fig. 3. For SPIs I and III, van der Waals binding interactions contribute more than the electrostatic interactions, whereas, for SPI-II, van der Waals and electrostatic binding energies are close to each other and contribute almost equally to the total binding. However, overall electrostatic interactions are reduced by the solvent effect as the electrostatic term of solvation free energies for binding (GB) are highly unfavorable. Nonpolar solvation free energies add to the favorable binding, but their contribution could be overestimated due to the lack of dispersion and repulsion terms in the ASP model.

**Fig. 3.**
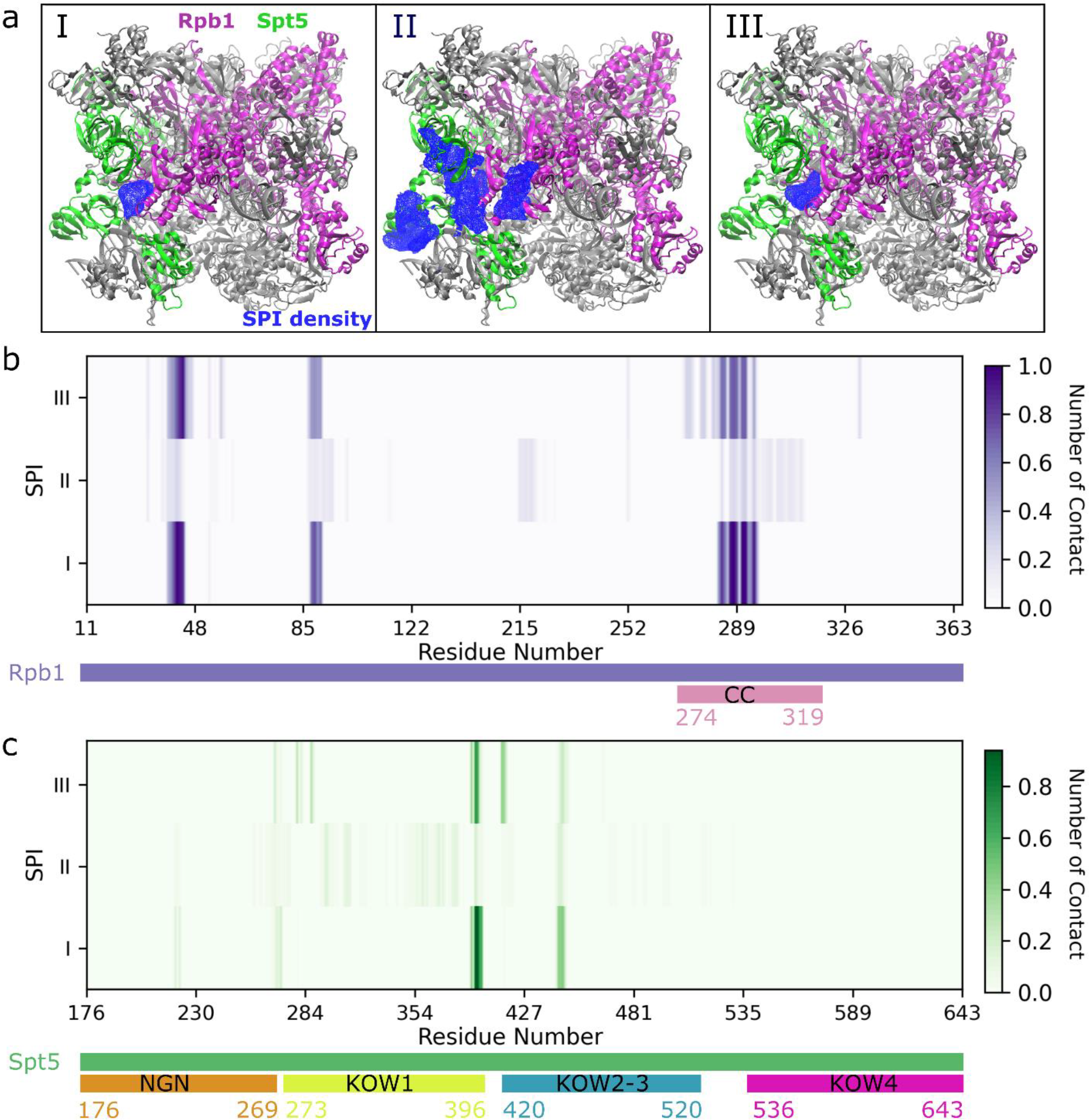
(a) The densities of SPIs were calculated for the trajectories of three replicates for I, II and III. Rpb1 is in magenta, Spt5 is in green, and the rest of the complex is in silver. SPI densities were shown in blue with 2 % iso-value occupancy. (b, c) Normalized number of contacts of SPIs with Rpb1 (b) and Spt5 (c). Rpb1 contacts are observed only for residues between 11-363. The CC domain of Rpb1 and NGN, KOW1, KOW2-3 and KOW4 domains of Spt5 were shown below the plots in different colors.

To obtain mostly populated binding sites and interaction networks, we applied PCA on the relative position of SPIs with respect to the center of mass of the Pol II-DSIF complex. Fig. 4a shows that I and III are located around the same region in the PCA plot as their binding sites are similar, while II spans a larger area in the PCA plot consistent with the densities observed in Fig. 3a. Fig. 4b shows the free energy plot obtained from the PCA, where we extracted the most populated (minimum energy) binding sites for each SPI shown in Fig. 4c, d, and e for I, II and III, respectively. We extracted one major binding position for I and III (Fig. 4c and e), and three positions for II (Fig. 4d). I_a_ has interactions with both Rpb1 and Spt5 (Fig. 4c), while III_a_ interacts mostly with Rpb1 residues rather than Spt5 (Fig. 4e). For I_a_, the aromatic rings are located inside a hydrophobic pocket made up by V284, Y43 and the hydrophobic chains of K42 and R291. On the other hand, III_a_ has more hydrophilic interactions; forms H-bonds with E89 and N288 and has electrostatic interactions with R292.

**Fig. 4.**
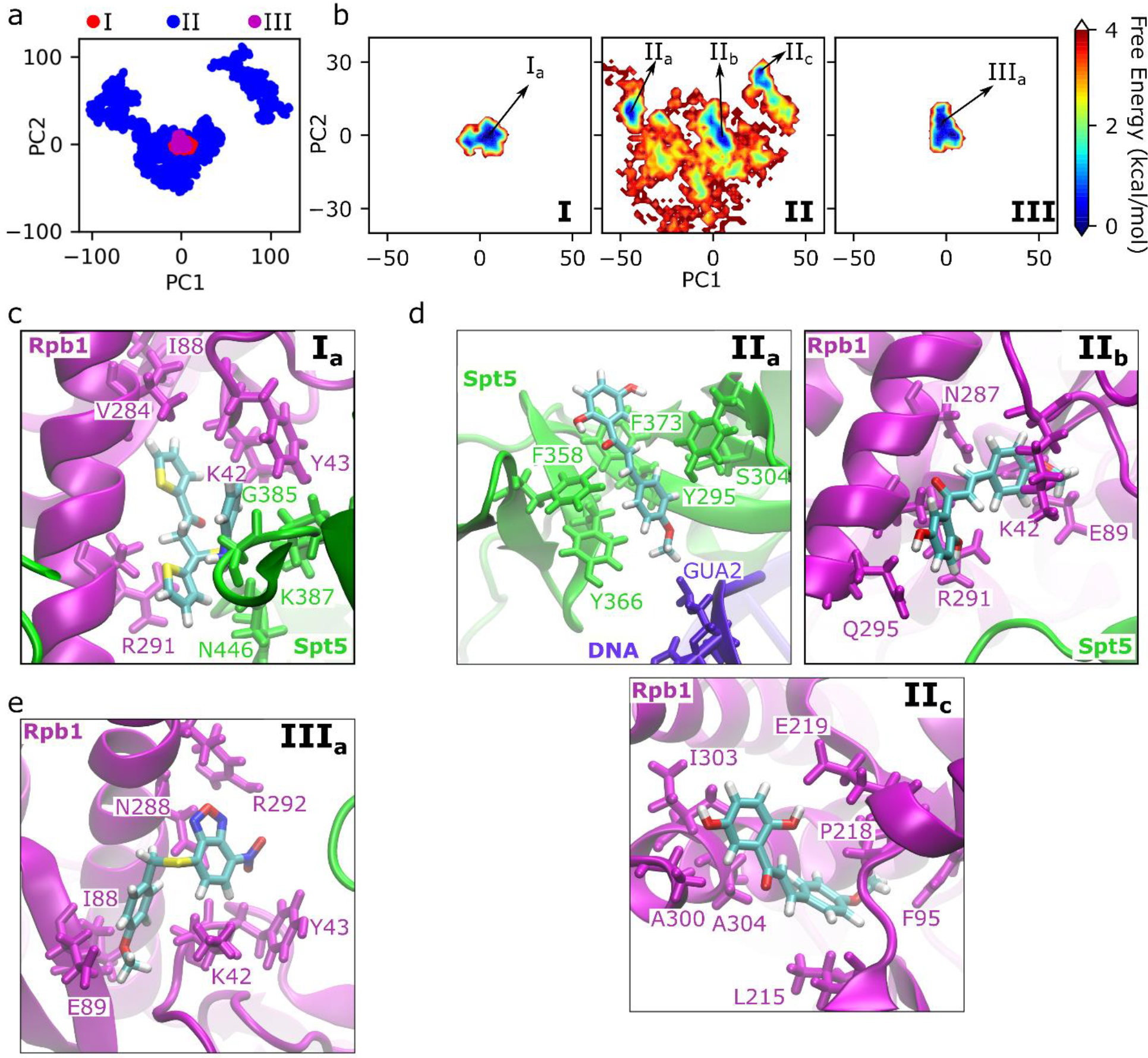
Binding positions of SPIs obtained from the PCA of the relative center of masses of SPIs with respect to the center of mass of the complete Pol II-DSIF complex. (a) PCA for three SPIs together; (b) potential mean force for the PC1 and PC2 separately calculated for three SPIs by applying WHAM analysis; (c-e) lowest energy positions obtained from the free energy plots for SPIs (c) I, (d) II, and (e) III.

For II, we extracted three positions, which are shown as II_a_, II_b_ and II_c_ in Fig. 4d. The II_a_ position was at the interface between the KOW1 domain of Spt5 and the upstream DNA without demonstrating any interactions with Pol II. It was located at the hydrophobic pocket formed by aromatic residues, which are F358, F373, Y295 and Y366 of Spt5. The II_b_ position is around the initial docking pose, where the molecule is located between the CC domain of Rpb1 and Spt5. It was surrounded by polar and charged residues which are stabilizing the SPI via H-bonds. The II_c_ position was also at the CC domain but facing the opposite side from the Spt5. It was stabilized by hydrophobic interaction with the residues I303, A300, A304, F95 and L215, and it also forms H-bond with E219. Overall, II was located around the CC domain and Spt5, but presented multiple binding positions in contrast to the single binding sites of I and III.

### SPIs impact DNA and RNA stability

To understand the effects of SPIs on the dynamics of Pol II, Spt5 and DNA/RNA at the exit tunnels, we analyzed RMSF and RMSD values of proteins and nucleic acids along the simulations. Fig. 5a shows the RMSF plots for the sections of the template and non-template DNA that are interacting with the KOW1 domain of Spt5 and the nascent RNA that is interacting with the KOW4 domain of Spt5. We observed increased fluctuations in DNA for the systems with SPIs I and III, while fluctuations were relatively smaller for the system with II compared to apo. As SPI-I and III were more strongly bound to the Pol II complex, this result suggests that the binding of SPIs to the Rpb1-Spt5 interface may destabilize the upstream DNA, which is known to be stabilized by interactions with Spt5 [7,16]. For the nascent RNA, we observed an opposite effect with reduced fluctuations in the RMSF plots for the systems with SPIs I, II and III. In addition to this, we observed high RMSD values for RNA as expected since it does not have a globular shape, while the RMSD values were relatively smaller and converged more quickly for the systems with I and III (Fig. 5b). On the other hand, RMSD values of DNA (Fig. S7) are mostly similar for the complexes with SPIs compared to the apo-complex, except that one replicate for each SPI-I and III showed larger RMSD values for DNA that supports a decreased stability of DNA observed in the RMSF plots.

**Fig. 5.**
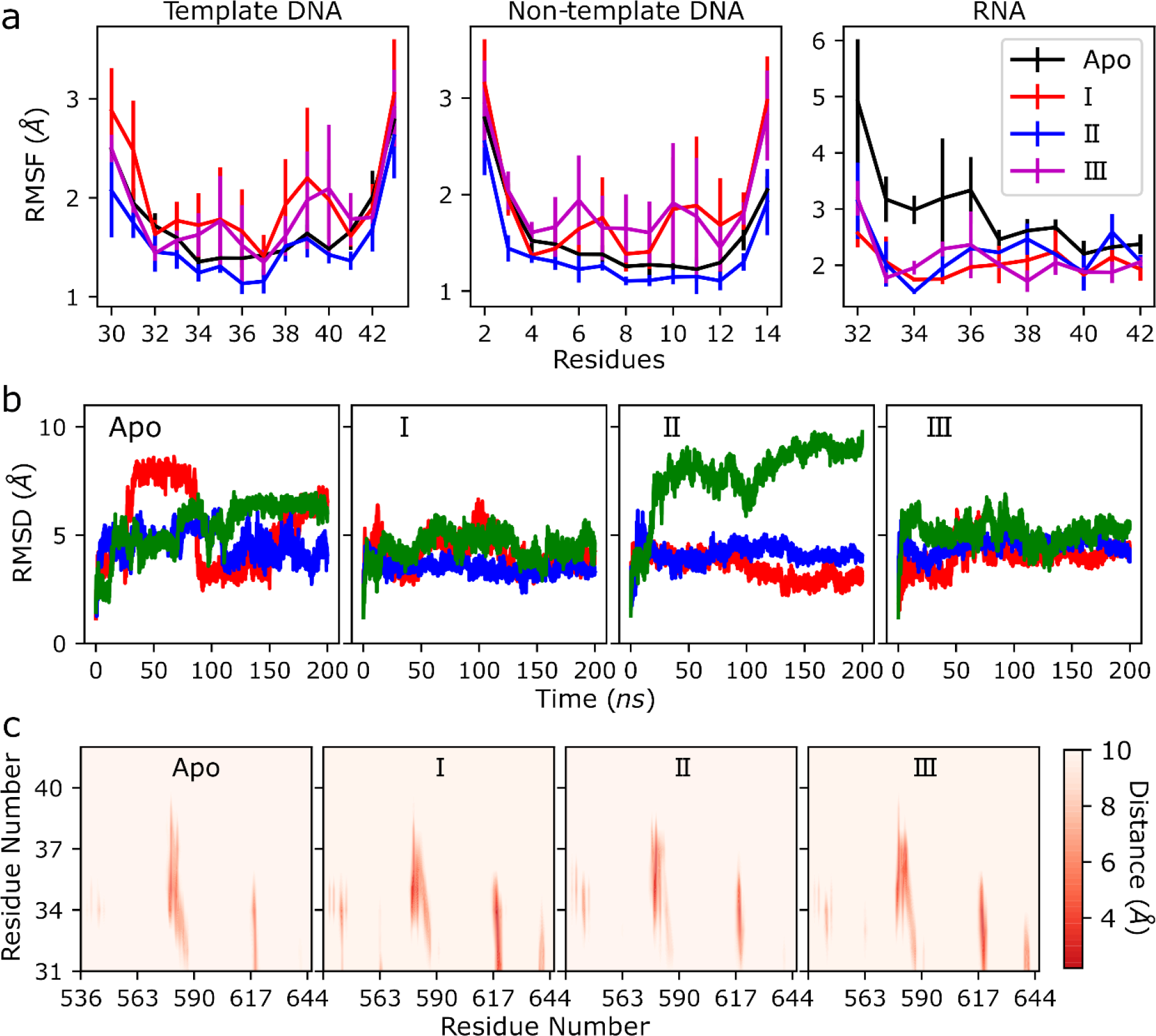
Analysis of the dynamics and interactions of upstream nucleic acids. (a) RMSF of template and non-template DNA and nascent RNA; (b) RMSD of the nascent RNA, red, blue and green lines represent the values of each replicate simulation; (c) distance maps between the KOW4 domain of Spt5 (x-axis) and the nascent RNA (y-axis).

To further understand the change in the stability of RNA, we focused on the interactions of RNA and the KOW4 domain of Spt5. Since the SPIs did not demonstrate any direct interactions with RNA, we assumed that SPIs impact the stability of RNA through the interactions of Spt5. Fig. 5c shows the distance maps of RNA with the KOW4 domain of Spt5. Apo showed strong interactions for KOW4 residues 578-585 and 617-620 and weaker interactions for 539-541. All these interaction sites were observed in the complexes with SPIs with reduced average distances suggesting the formation of stronger interactions compared to the apo-protein. Furthermore, additional interaction sites (563-564 and 639-642) were observed for systems with I and III. The analyses of the distance maps and RMSF/RMSD of RNA suggested that the SPIs increased the interactions between Spt5 and RNA and, as a result, stabilized RNA at the exit tunnel. For DNA, Figs. S8 and S9 show the distance maps between the KOW1 domain of Spt5 and non-template and template DNA at the exit tunnel. There are not any significant changes in the distance maps in the presence of SPIs, while interactions with the template chain somewhat weakened especially for the residues between 361-366.

For the proteins, we calculated RMSF values of the CC domain of Rpb1, and NGN, KOW1 and KOW4 domains of Spt5 (Fig. S10). For the CC, we observed some small changes in the flexible parts of the domain (residues 290-302 and 320-326). For Spt5, there are not any significant changes in the RMSF values, except for the increased fluctuations at the KOW1 domain (residues 344-357) with SPI III. RMSD values were slightly higher for the apo-complex and the complex with II for Spt5 (Fig. S11), while RMSD values for Rpb1 did not show any significant changes (Fig. S12).

Overall, RMSF and RMSD plots of the nascent RNA suggest that the SPIs, especially I and III, cause stabilization of RNA at the exit tunnel; and distance maps suggest that SPIs I and III indirectly impact RNA dynamics by increasing interactions between the Spt5-KOW4 domain and RNA. On the other hand, upstream DNA was destabilized in the presence of I and III, and interactions between Spt5-KOW1 and DNA were slightly weakened suggesting an indirect impact of SPIs on DNA dynamics as well, but in the opposite direction. To understand the different effects of SPIs on DNA and RNA dynamics, we focused on SPI-I and performed PCA on the distances between the Spt5-KOW4 domain and RNA, since they showed larger differences between the SPI-systems and apo-complex compared to KOW1-DNA interactions. Fig. 6a shows that SPI-I system covers a smaller space than the apo-complex, suggesting that KOW4-RNA distances fluctuate less in the presence of SPI-I. This is consistent with the reduced dynamics of RNA observed in RMSF plots shown in Fig. 5a. We extracted the minimum energy structures from the PCA free energy plot. Fig. 6b shows the location of SPI-I, which is close to KOW1 and KOW2-3 domains, while far from the KOW4 domain of Spt5 consistent with the contact map analysis shown in Fig. 3c. SPI-I is also not directly in contact either with RNA or DNA at the exit tunnels, therefore its effects on the dynamics of RNA and DNA are expected to be indirect. Fig. 6c suggests that in the presence of SPI-I, KOW1 moved slightly away from the DNA exit potentially due to the interactions with the SPI-I, together with the DNA partly lost interactions with Spt4 (Fig. S13) and probably as a result, DNA exit channel opened more, which then resulted in higher dynamics of DNA compared to the apo-complex. On the other hand, Fig. 6d shows an opposite trend in the RNA dynamics as RNA exit channel became tighter in the presence of SPI-I. RNA exit site is formed by the Spt5-KOW4, Rpb1 dock and Rpb2 wall domains. In the presence of SPI-I the interactions of wall domain with KOW1 and KOW4 increase (Fig. S14). These increased interactions made the RNA exit tunnel tighter, and potentially caused the further stabilization of RNA.

**Fig. 6.**
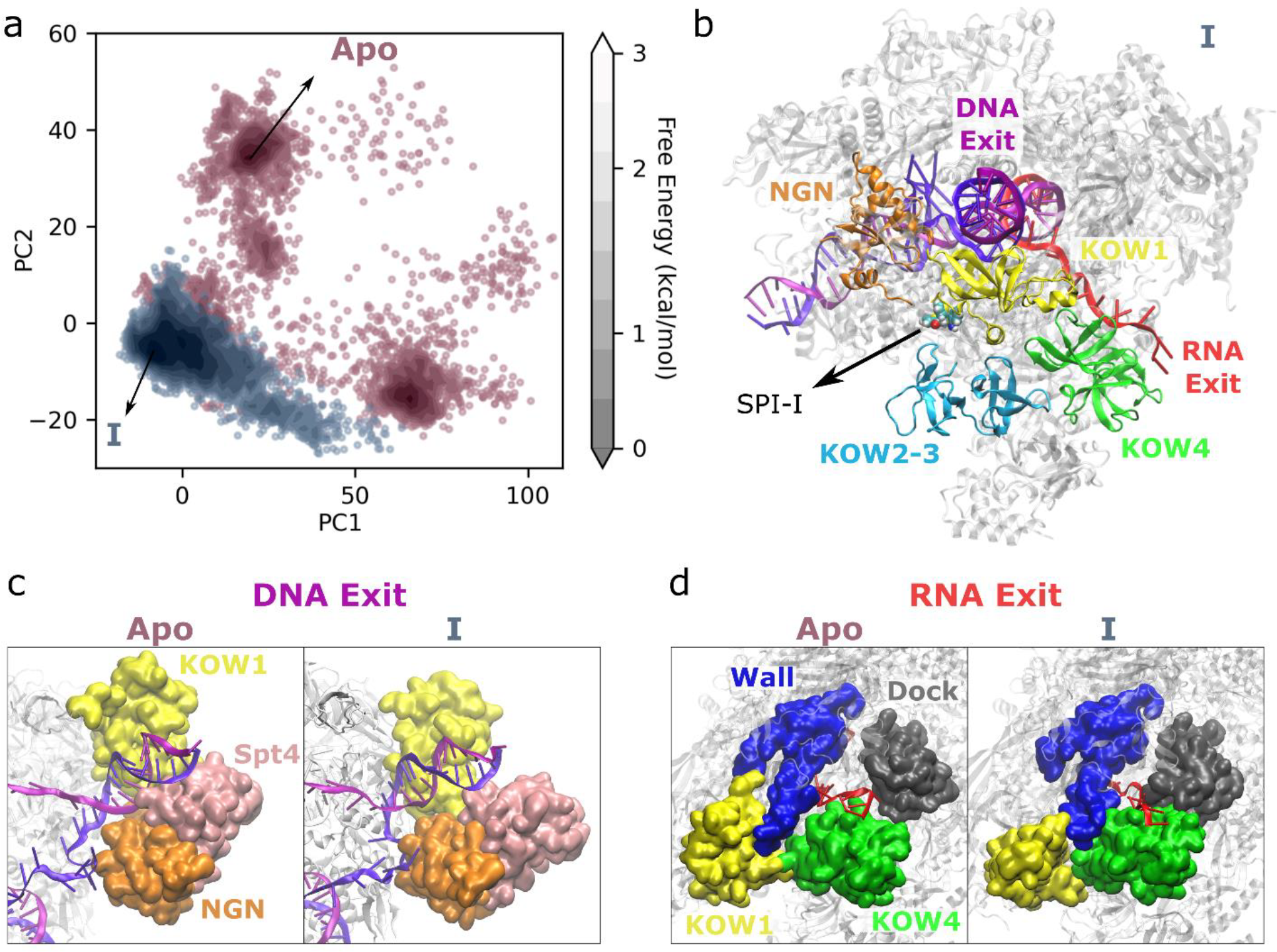
Analysis of the apo-complex and the complex with SPI-I based on RNA-KOW4 distances. (a) PCA plot that shows the free enery contour plots in gray scale and each point from apo-complex and the complex with SPI-I shown in red and blue, respectively; (b) the minimum energy structure of the complete complex with SPI-I extracted from the simulation trajectories based on the PCA plot, the NGN, KOW1, KOW2-3, KOW4 domains of Spt5 are shown in different colors, SPI-I is shown in vdW representation, RNA and DNA exits are labeled in red and magenta, respectively; DNA (c) and RNA (d) exit sites from minimum energy structures of the apo-complex and the complex with the SPI-I extracted from the simulation trajectories based on the PCA plot.

## DISCUSSION

In this study, we predicted binding sites of potential drug molecules that are known to inhibit the mutant Htt gene transcription. We studied three SPIs, I, II and III, that were reported earlier by Bahat et al with the numbers 18, 21 and 86. We choose I and II since there are experimental measurements for these two SPIs that suggest direct interactions between them and Rpb1 and/or Spt5, therefore, we can compare the experimental results with our binding predictions. Experiments suggested that both SPI-I and II have direct interaction with the Rpb1 CC domain and SPI-I also has interactions with Spt5. We applied docking to a grid that covers these experimentally predicted interaction sites rather than using a large grid to cover the complete protein. We applied docking to a cryo-EM Pol II-DSIF structure, which has relatively low resolution (3.7 Å) [16]. Studies suggest that accuracies of docking decreases with decrease in resolution (lower resolution than 2.5 Å) [59-61]. In addition to this, using large grid sizes are known to reduce the accuracies of docking and potentially provide irrelevant binding poses [62]. Therefore it is particularly challenging to apply docking on Pol II-DSIF complex and we decided to use experimental bias in our docking procedure to overcome these limitations.

Binding of SPI-I and II to the Pol II-DSIF complex was monitored experimentally using fluorescence intensity measurements by Bahat *et al*. [36]. Experiments showed that SPI-I affected intrinsic fluoresence of Spt5 and Rpb1 while II showed effect only on the fluorescence spectrum of Rpb1 suggesting that SPI-I has direct interactions with both Rpb1 and Spt5, while SPI-II has interactions only with Rpb1. In addition to these binding assay results, the measurement of half-maximal inhibitory concentrations (IC_50_) in enzymatic acitivity assays show that SPI-I, II and III have IC_50_ values of 33.1, 25.2, and 5.5 μM, respectively, suggesting that they are biologically active and these IC_50_ values are comparable to the IC_50_ values obtained by fluorescence intensity measurements which suggest that their activity might be directly related to their binding to Rpb1 and Spt5. During the MD simulations, consistent with experiments, we observed that SPI-I was located at the interface of Rpb1 and Spt5 and had interactions with both the CC-domain and Spt5, while SPI-II manifested multiple binding sites that include both the CC domain of Rpb1 and Spt5. Additionally, SPI-III also shared a similar binding site with SPI-I, located at the interface of Rpb1 and Spt5. Binding to the Pol II-DSIF complex was stronger for I and III, compared to II and this may be related to their effects on the inhibition of the mutant Htt gene transcription. SPI I and II both selectively inhibited mutant Htt gene transcription, but SPI-II had significant effects on the transcription of inflammatory genes at both basal and induced conditions while SPI-I had only effects at induced conditions [36]. SPI-III has inhibition patterns similar to II, but shows stronger inhibition of the mutant Htt gene transcription as SPI-III inhibited it at very small concentrations; reported 0.1 μM compared to 30 μM of I and II [36]. The inhibition of transcription of a wider range of genes by SPI-II could be related to its multiple binding sites, which may cause multiple mechanisms of inhibiting different genes including inflammatory genes and Htt gene. However, this hypothesis needs to be further tested by experimental and computational studies.

We calculated the binding energies of SPIs from the MD simulation trajectories using MM/GBSA method. The binding energies suggested more favorable interactions for SPI-I and III compared to II, which is consistent with the strong binding to the initital binding sites for SPI-I and III, while a relatively loose binding led SPI II to sample multiple binding sites. Although the trend of binding enegies matched with the observed behaviour during the simulations, the experimental IC_50_ values from binding assays suggest that SPI-II more strongly binds to Pol II than SPI-I. Similarly, SPI-II has a smaller IC_50_ in the enzymatic activity assays than SPI-I [36]. On the other hand, the most favorable binding was observed for SPI-III in MM/GBSA calculations and the experimental IC_50_ values from the ezymatic actitivity assays of SPI-III is the smallest among the three SPIs, which support the strong binding for this SPI. We also note that MM/GBSA method is known to overestimate binding energies toward more favorable binding [63]. Although it is difficult to assess the overestimation for our case as there are not any available experimental binding energies for the SPIs, we obtained highly favorable energies that may be unrealistic. The MM/GBSA method has many limitations that may result in overestimation of the binding. One limitation is coming from the calculation of the electrostatic solvation free energy by the GB formula that approximates the dielectric constant of solute as 1. However, studies suggest that the dielectric constant depends on the binding site especially for charged or polar residues and using 1 regardless of the solute characteristics may cause deviations of calculated binding energies from the experimental results [63-65]. Another limitation is coming from the calculation of the nonpolar solvation free energy, which includes only the cavity term in our calculation. The lack of the dispersion and repulsion energy terms may cause deviations in the energy calculations. Although the contribution from the dispersion and repulsion terms are relatively small, it could make a difference especially for the hydrophobic sites. In addition to this, the cavity term calculated by SASA does not account for the water molecules in the cavity of the binding site in the absence of the ligand as it uses continuum solvent models. Therefore, calculating the nonpolar solvation free energy with SASA model may cause additional deviations from the actual binding energies [63,66]. Lastly, in our MM/GBSA calculations, we excluded the entropy term as the calculation of this term is computationally expensive especially for a large protein as Pol II. There could be an entropic barrier for the SPIs to change the conformations upon binding to the protein, which could potentially make the binding of the ligands less favorable.

We observed that SPIs affect the dynamics of RNA and DNA at the exit tunnel. The effects are larger for SPIs I and III, as expected since they demonstrated a stronger binding than II. We showed that Spt5-RNA interactions became stronger in the presence of SPIs suggesting that SPIs indirectly affect RNA dynamics through interactions with Spt5. Similarly, Spt5-DNA interactions became weaker, and this potentially caused the increased flexibility of the DNA chains, however the impact of SPIs on Spt5-DNA interactions was weaker compared to Spt5-RNA interactions. Overall, our observations suggest that SPIs stabilize RNA and destabilize DNA. However, in this study, we directly used the DNA template sequence and the sequence of the corresponding nascent RNA that were present in the transcription bubble of the initial cryo-EM structure rather than using a DNA/RNA sequence with CAG repeats that is relevant to Htt gene. The study from Bahat *et al*. [36] suggested that SPIs selectively affect the transcription of the long repeat Htt genes rather than short repeat genes. The earlier studies also showed that Spt4/Spt5 play an important role in the mutant Htt gene transcription selectively [12,15]. The findings from these studies suggest that long repeat genes increases the requirement of DSIF for transcription and, thus, interfering in the DSIF function selectively inhibits the transcription of these genes. We hypothesize that the increased function of DSIF in regulating the long repeat Htt gene transcription is potentially related to the conformational changes in the Pol II-DSIF complex in the presence of the long repeat genes. Therefore, to obtain a mechanistic understanding of the inhibition of mutant Htt gene transcription, we need to perform simulations with the mutant Htt gene in the transcription bubble. There is not any available structure of Pol II-DSIF complex with the mutant or wild type (WT) Htt genes. Therefore, performing additional simulations with Htt genes requires an extensive modeling of the DNA-RNA hybrid in the transcription bubble by mutating the nucleic acids in the available structures and potentially utilizing the published structures of RNA and DNA CAG repeats [67-69]. However, we expect large conformational changes in the complex in the presence of long repeat genes and to capture such conformational changes, an exhaustive modeling followed by μs long extensive MD simulations would be required. We expect that RNA stabilization at the exit site could be larger with mutant Htt gene that may induce pausing, or alternatively largely destabilized upstream DNA may cause transcription defects.

Lastly, we note that stereochemistry of the SPIs may also have effects in their inhibition activity. SPI-I has two enantiomers, which have different binding patterns according to our simulations. R-I that is presented in the main text, shows a strong binding to the initial binding site, while S-I sampled larger number of binding sites, mostly around Rpb1 (Fig. S2). Their effects on nucleic acid dynamics also varied as they both stabilized RNA, but R-I increased the fluctuations for the DNA conformations while S-I showed smaller impact on the dynamics of DNA. These results suggest that R-I could be more active enantiomer than S-I or alternatively their inhibition mechanisms and which genes they inhibit may be distinct. Future experimental studies would help to understand the differences in the inhibition mechanisms of these two enantiomers.

## CONCLUSION

In this study, we propose feasible binding modes to the Pol II-DSIF complex for three recently discovered SPIs. We observed that two out of three SPIs were strongly bound to the Spt5 and Pol II interface as suggested by experiments, while the remaining SPI bound to multiple binding sites. SPIs indirectly affected dynamics of the nascent RNA and upstream DNA via interactions with Spt5. We concluded that the transcription inhibition mechanism could be related to Spt5-nucleic acid interactions. Future work will focus on computational modeling and the simulation of the Pol II elongation complex bound to the relevant regions of WT and mutant Htt genes to investigate further the role of Spt5 in the inhibition mechanisms of multi-repeat genes.

## Acknowledgments

We used the computational resources at the MERCED and Pinnacles clusters at the University of California Merced and the Advanced Cyberinfrastructure Coordination Ecosystem: Services & Support (ACCESS) facilities funded by the National Science Foundation (ACCESS award number is TG-BIO210145). We thank Brandon Michael Bogart for setting up the initial systems for the apo elongation complex simulations.

## Author contributions

*Adan Gallardo:* Conceptualization, Formal Analysis and Investigation, Methodology, Visualization, Writing – Original Draft Preparation, Writing – Review & Editing.

*Bercem Dutagaci:* Conceptualization, Formal Analysis and Investigation, Methodology, Visualization, Writing – Original Draft Preparation, Writing – Review & Editing, Project Administration, Resources, Supervision.

## Supporting Information

Figures S1-S14 are provided as Supporting Information.

## Data availability statement

The initial structure of the Pol II elongation complex was obtained from the Protein Data Bank with the PDB ID:5OIK. Details of the generation of initial SPI structures and their complexes with Pol II elongation complex were provided in the Methods section. We used open software for the simulations (OpenMM) and analysis (MDAnalysis and MMTSB). We provided the extracted frames from the PCA plots on our GitHub page (https://github.com/bercemd/SPIs_binding). Simulation trajectories can be available upon request.

