## Supplementary Information for "Binding of Small Molecule Inhibitors to RNA Polymerase-Spt5 Complex Impacts RNA and DNA Stability"

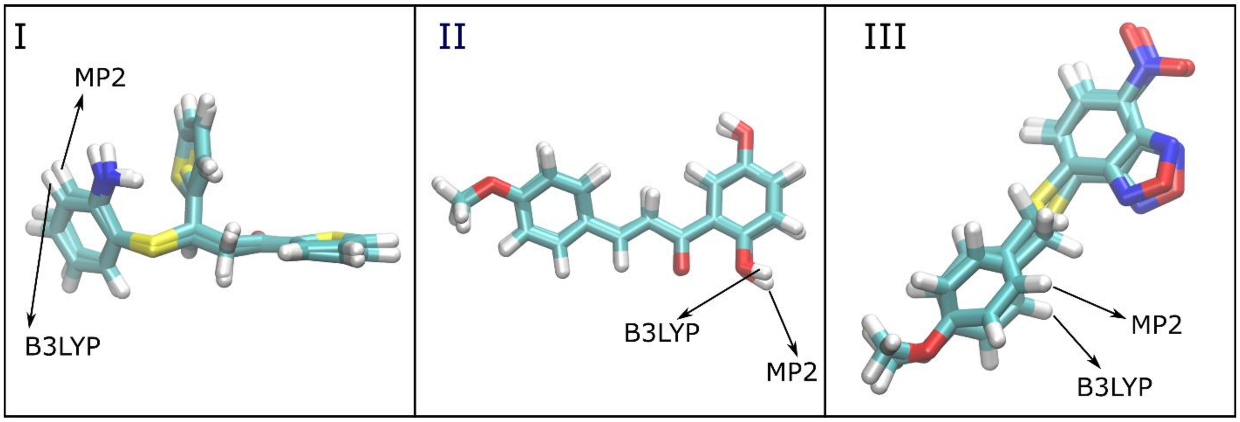


**Figure S1.** 3D structures of SPIs obtained by geometry optimization using the MP2 and B3LYP algorithms.

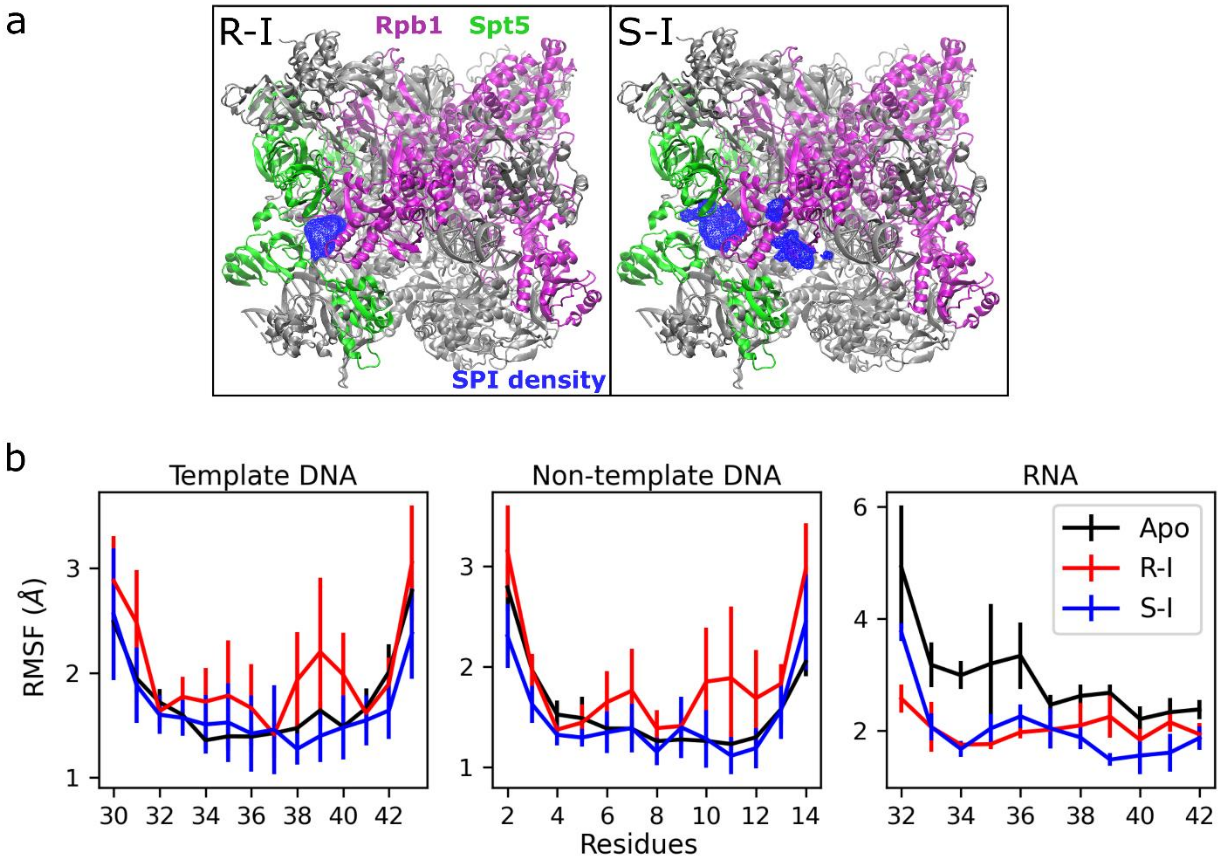


**Figure S2.** (a) The densities of SPIs were calculated for the trajectories of three replicates for R (R-I) and S (S-I) enantiomers of SPI-I. Rpb1 is in magenta, Spt5 is in green, and the rest of the complex is in silver. SPI densities were shown in blue with 2 % iso-value occupancy. (b) RMSF of template and non-template DNA and nascent RNA for the apo-complex, and the complexes with R-I and S-I.

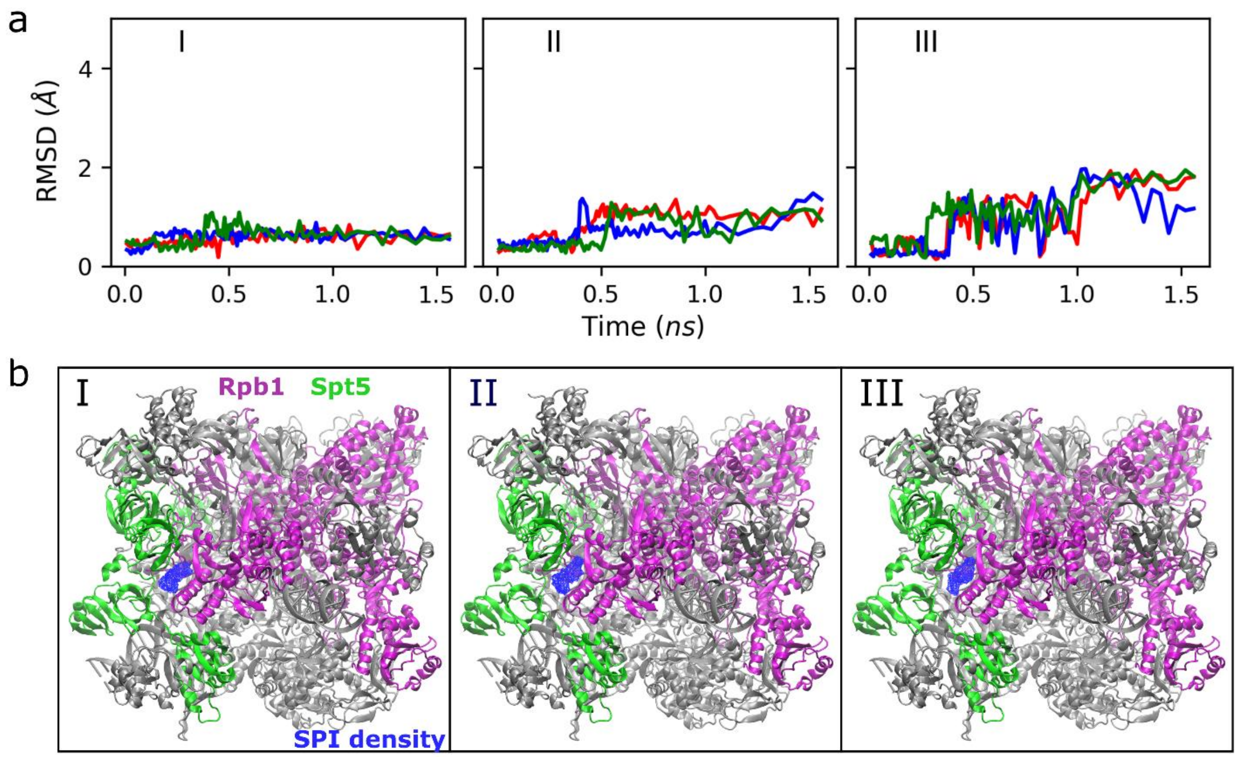


**Figure S3.** (a) RMSD values of SPIs during the equilibration; red, blue, and green lines represent the values of each replicate simulation. (b) The densities of SPIs were calculated for the equilibration trajectories of three replicates for I, II, and III. Rpb1 is in magenta, Spt5 is in green, and the rest of the complex is in silver. SPI densities were shown in blue with 2 % iso-value occupancy.

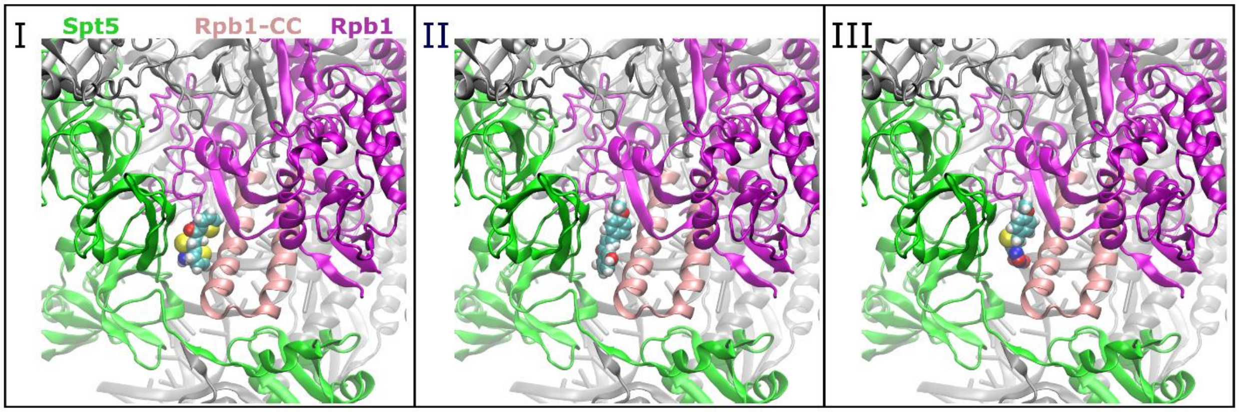


**Figure S4.** Docking positions for SPIs I, II, and III at the Pol II-DSIF complex. The color code for the Pol II-DSIF complex is as follows: Rpb1 is in magenta, the coiled-coil (CC) domain of Rpb1 is in pink, Spt5 is in green, and the rest of the complex is in silver. The SPIs are shown in colors coded by atom name: C is cyan, H is white, O is red, N is blue, and S is yellow.

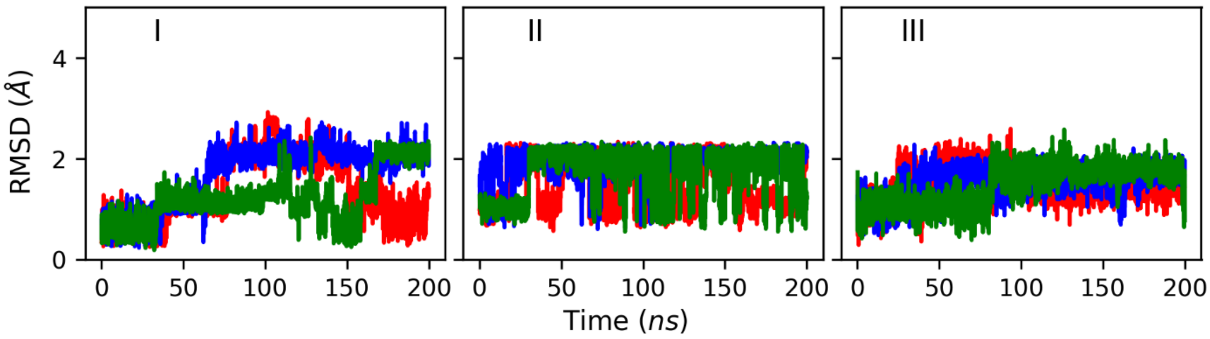


**Figure S5.** RMSD values of SPIs. Red, blue, and green lines represent the values of each replicate simulation.

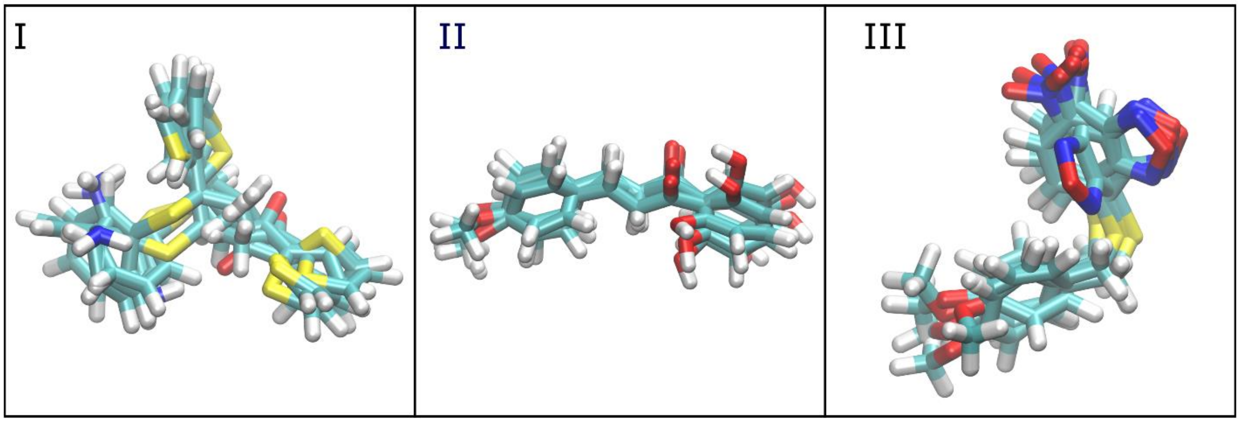


**Figure S6.** Central structures of SPIs from the five clusters obtained by Kmeans clustering algorithm based on RMSD values along the MD simulations.

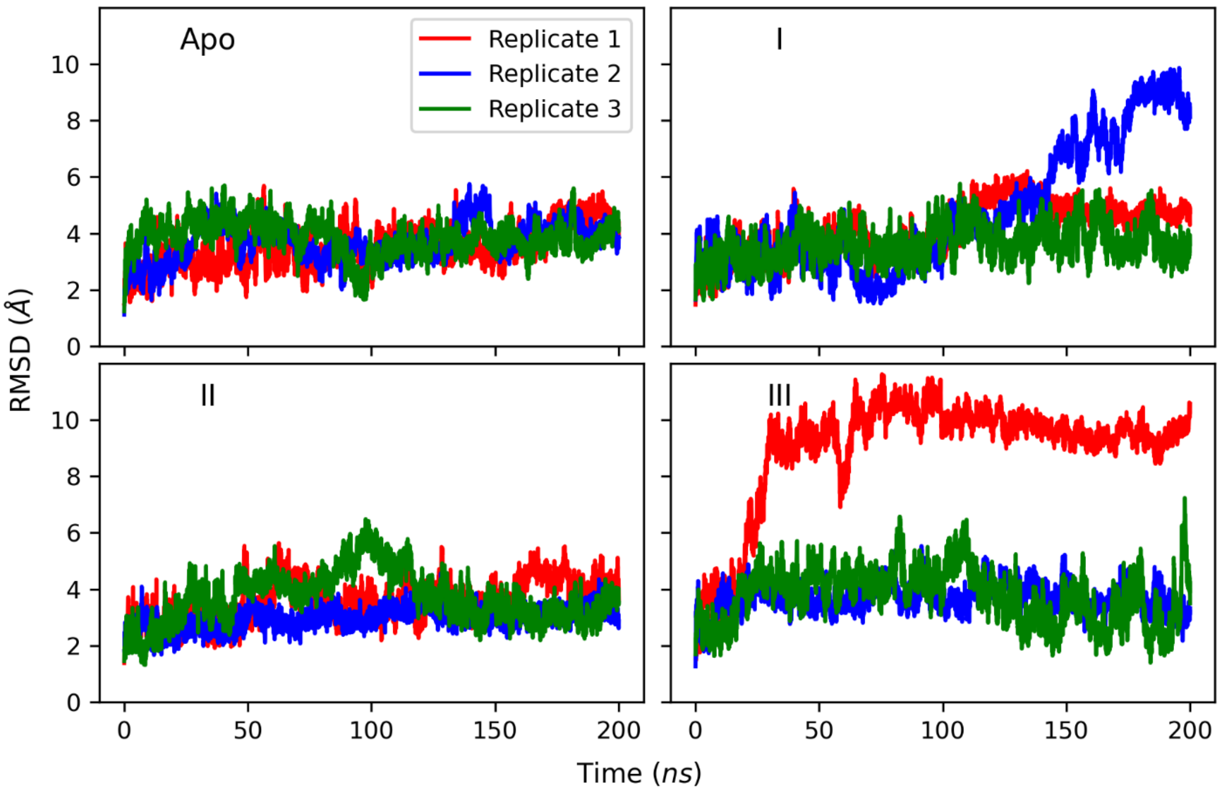


**Figure S7.** RMSD values of upstream DNA. RMSDs were calculated for the combined residues of template (residue numbers 30-43) and non-template (1-14) DNA chains. Red, blue, and green lines represent the values of each replicate simulation.

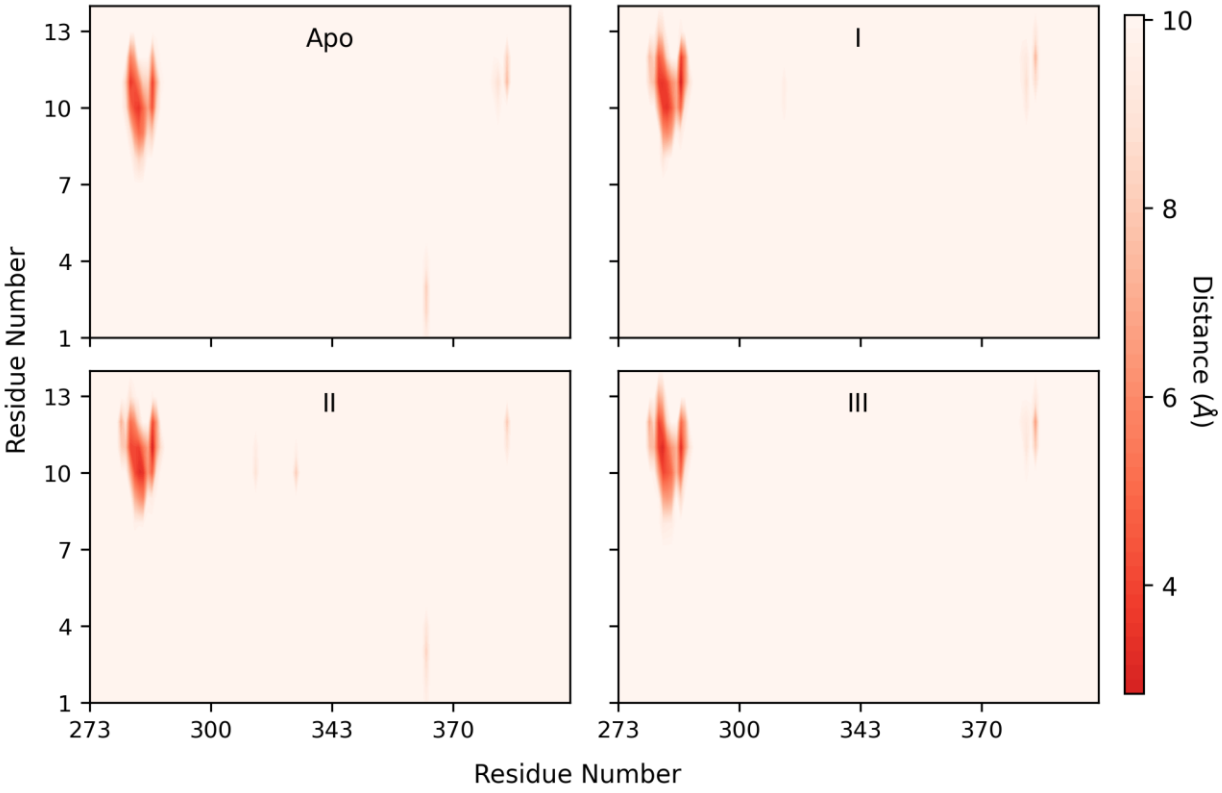


**Figure S8.** Distance maps between the KOW1 domain of Spt5 (x-axis) and upstream non-template DNA (y-axis).

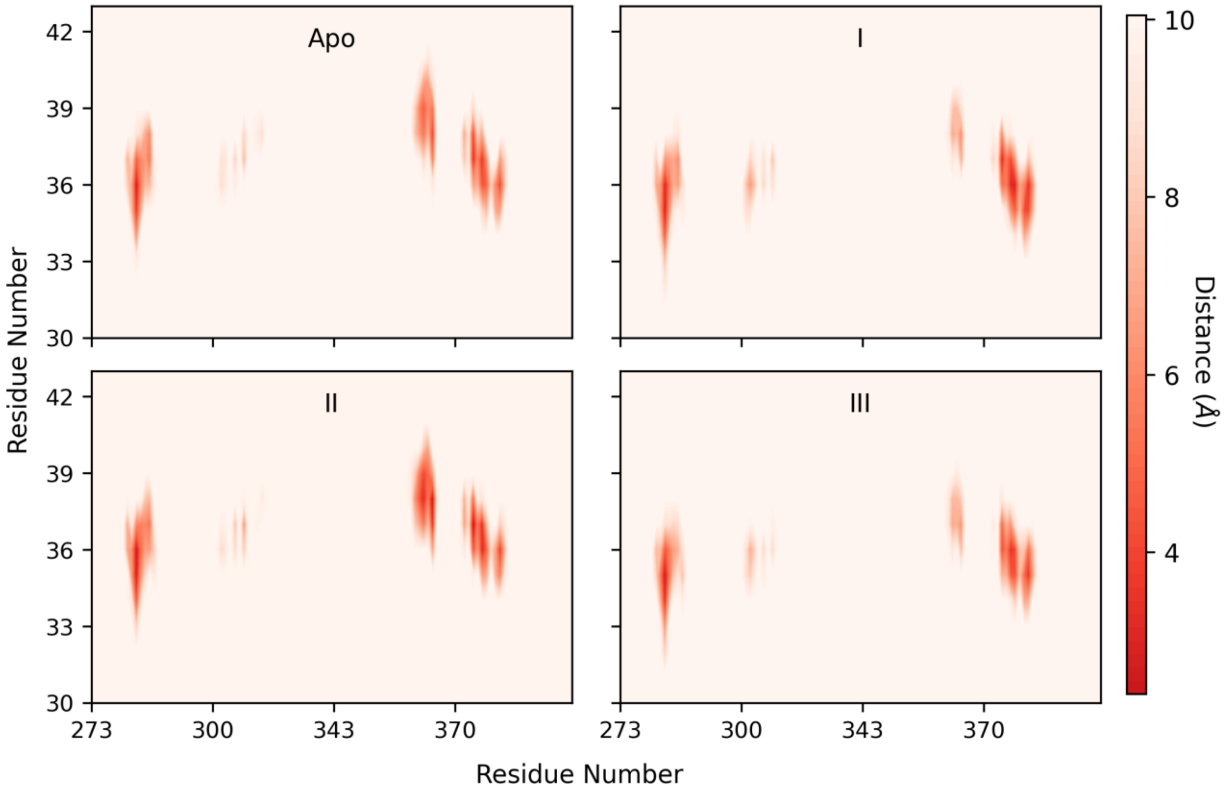


**Figure S9.** Distance maps between the KOW1 domain of Spt5 (x-axis) and upstream template DNA (y-axis).

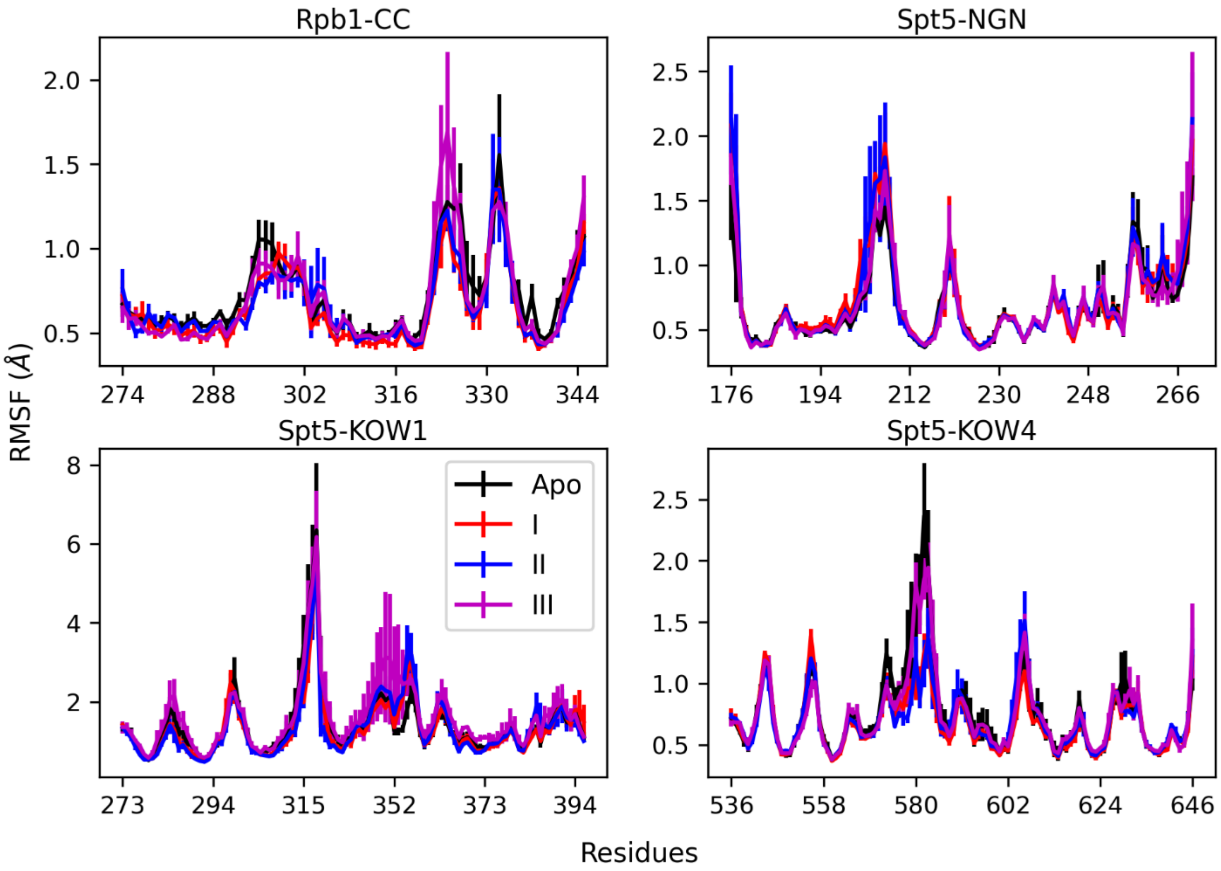


**Figure S10.** RMSF of the CC domain of Rpb1 and NGN, KOW1, and KOW4 domains of Spt5.

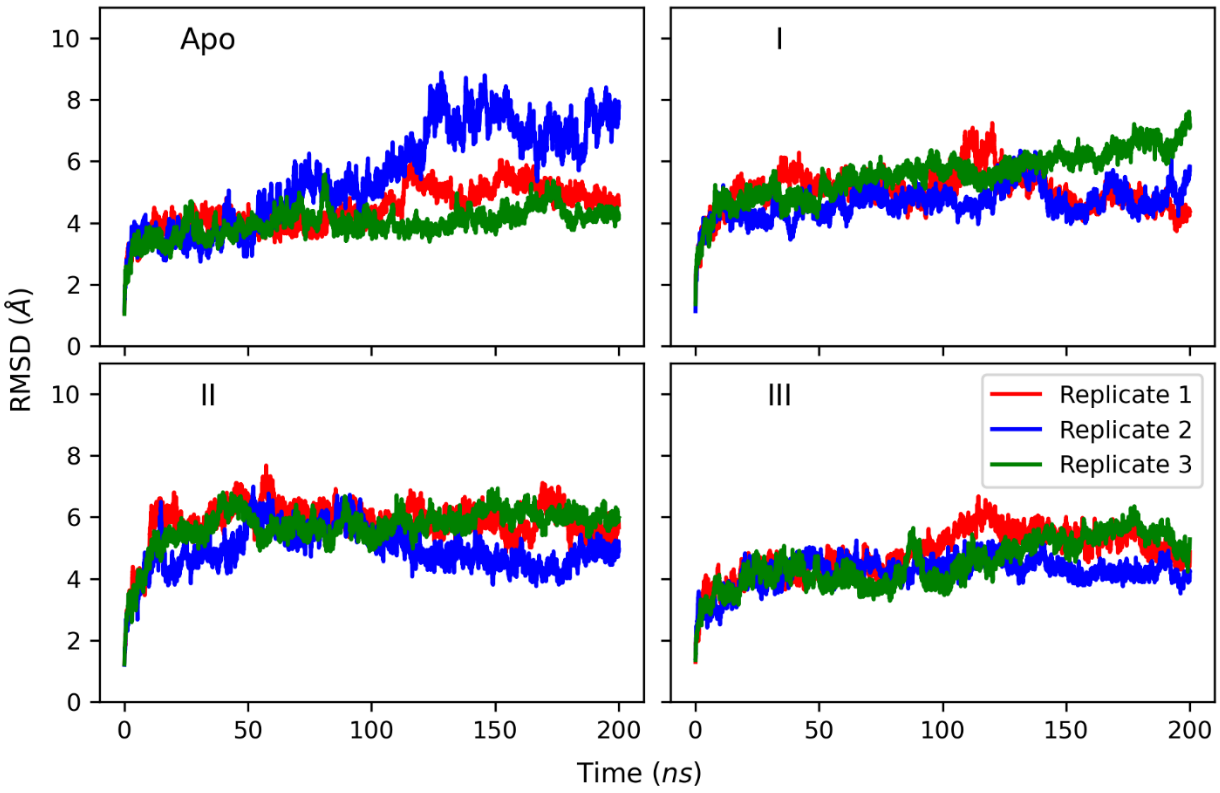


**Figure S11.** RMSD values of Spt5. Red, blue, and green lines represent the values of each replicate simulation.

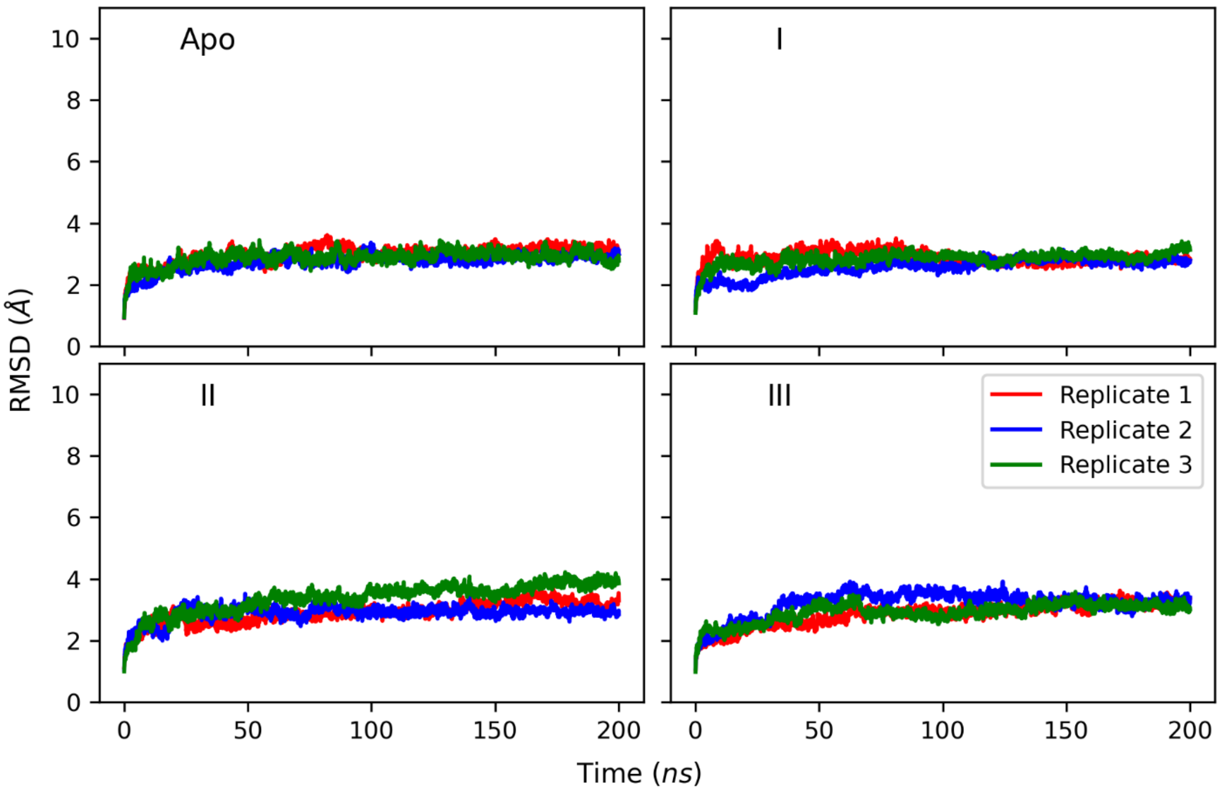


**Figure S12.** RMSD values of the Rpb1 subunit of Pol II. Red, blue, and green lines represent the values of each replicate simulation.

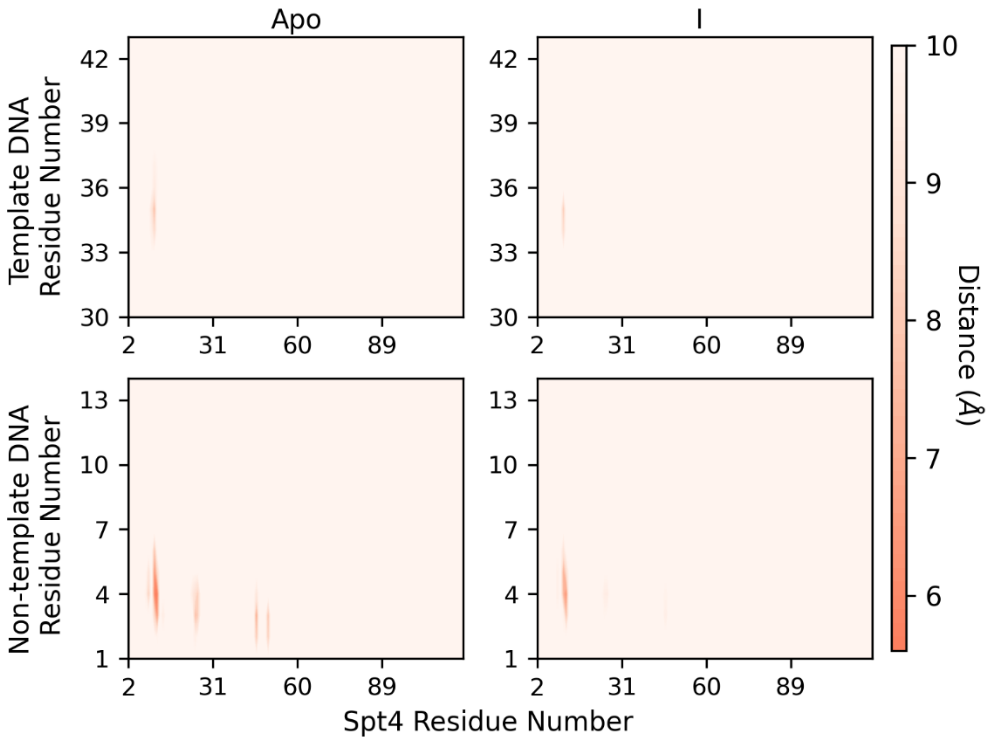


**Figure S13.** Distance maps between Spt4 and upstream template and non-template DNA for apo-complex and the complex with SPI-I.

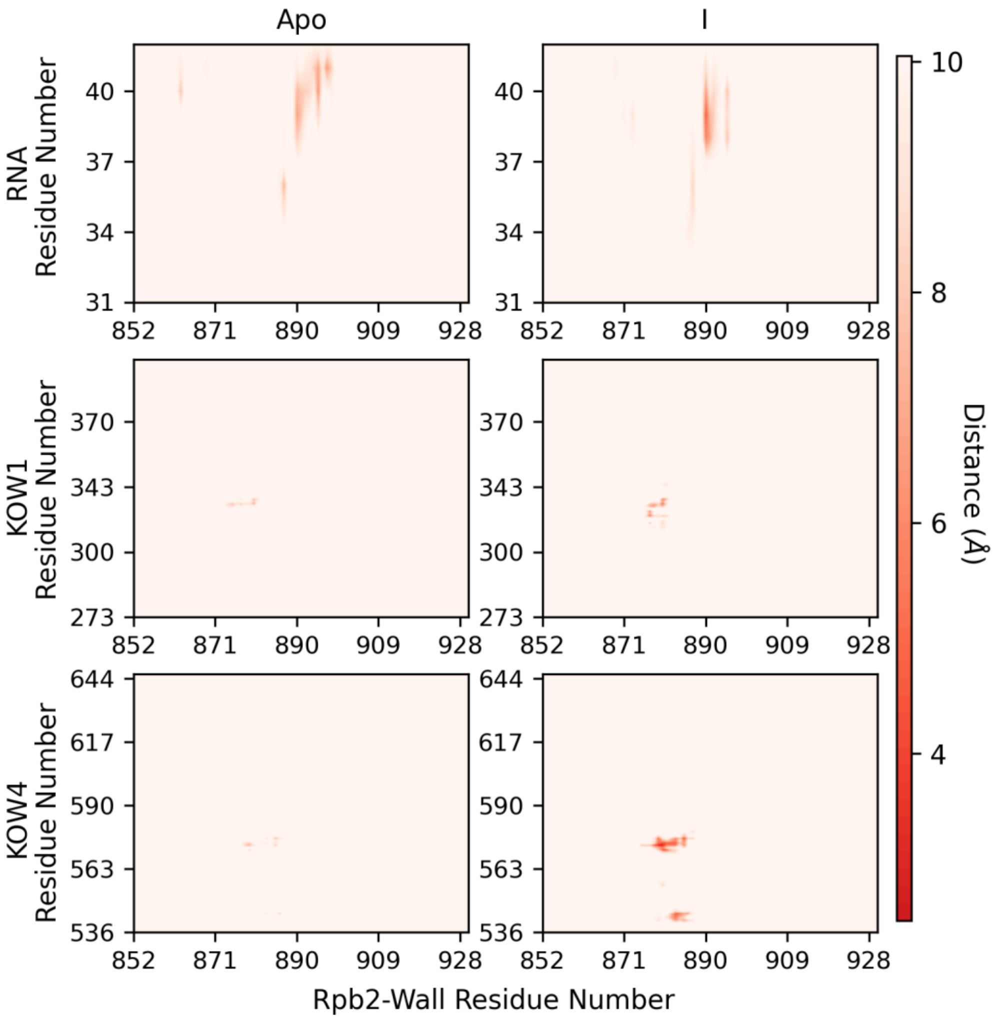


**Figure S14.** Distance maps between Rpb2 wall domain and RNA, KOW1, and KOW4 for apo-complex and the complex with SPI-I.
